# Benchmarking topological accuracy of bacterial phylogenomic workflows using *in silico* evolution

**DOI:** 10.1101/2021.08.03.454900

**Authors:** Boas C.L. van der Putten, Niek A.H. Huijsmans, Daniel R. Mende, Constance Schultsz

## Abstract

Phylogenetic analyses are widely used in microbiological research, for example to trace the progression of bacterial outbreaks based on whole-genome sequencing data. In practice, multiple analysis steps such as *de novo* assembly, alignment and phylogenetic inference are combined to form phylogenetic workflows. Comprehensive benchmarking of the accuracy of complete phylogenetic workflows is lacking.

To benchmark different phylogenetic workflows, we simulated bacterial evolution under a wide range of evolutionary models, varying the relative rates of substitution, insertion, deletion, gene duplication, gene loss and lateral gene transfer events. The generated datasets corresponded to a genetic diversity usually observed within bacterial species (≥95% average nucleotide identity). We replicated each simulation three times to assess replicability. In total, we benchmarked seventeen distinct phylogenetic workflows using 8 different simulated datasets.

We found that recently developed k-mer alignment methods such as kSNP and SKA achieve similar accuracy as reference mapping. The high accuracy of k-mer alignment methods can be explained by the large fractions of genomes these methods can align, relative to other approaches. We also found that the choice of *de novo* assembly algorithm influences the accuracy of phylogenetic reconstruction, with workflows employing SPAdes or SKESA outperforming those employing Velvet. Finally, we found that the results of phylogenetic benchmarking are highly variable between replicates.

We conclude that for phylogenomic reconstruction k-mer alignment methods are relevant alternatives to reference mapping at species level, especially in the absence of suitable reference genomes. We show *de novo* genome assembly accuracy to be an underappreciated parameter required for accurate phylogenomic reconstruction.

**Impact statement:** Phylogenetic analyses are crucial to understand the evolution and spread of microbes. Among their many applications is the reconstruction of transmission events which can provide information on the progression of pathogen outbreaks. For example, to investigate foodborne outbreaks such as the 2011 outbreak of *Escherichia coli* O104:H4 across Europe. As different microbes evolve differently, it is important to know which phylogenetic workflows are most accurate when working with diverse bacterial data. However, benchmarks usually consider only a limited dataset. We therefore employed a range of simulated evolutionary scenarios and benchmarked seventeen phylogenetic workflows on these simulated datasets. An advantage of our simulation approach is that we know *a priori* what the outcome of the analyses should be, allowing us to benchmark accuracy. We found significant differences between phylogenetic workflows and were able to dissect which factors contribute to phylogenetic analysis accuracy. Taken together, this new information will hopefully enable more accurate phylogenetic analysis of bacterial outbreaks.

**Data summary:** A Zenodo repository is available at https://doi.org/10.5281/zenodo.5036179 containing all simulated genomes, all alignments produced by phylogenetic workflows and .csv files summarising the topological accuracies of phylogenies produced based on these alignments. Code is available at https://github.com/niekh-13/phylogenetic_workflows.

## Introduction

Phylogenetic analyses are crucial to assess the relatedness within a population of micro-organisms. These analyses provide information on the speciation, evolution and spread of microbes. Within clinical settings they can be used to identify microbial outbreaks and transmission events^1^. With the introduction of cost-efficient whole-genome sequencing (WGS), bacterial outbreak tracing is increasingly based on whole genome data, instead of on a small section of the genome such as 16S rRNA genes or a set of universal genes^2^. Whole-genome phylogenetic analysis can be applied by various pipelines or workflows, often composed of multiple separate tools. Common differences between workflows are which genomic loci are considered in the analysis (only protein-coding genes or also intergenic regions), how genetic features are defined (genes, k-mers, single nucleotide variants, etc.), but also how genomes are assembled. Benchmarking is necessary to make sense out of the plethora of bioinformatic methodologies available. However, most benchmarks focus on a single or few datasets^3–5^, limiting the generalizability of their findings. Additionally, it is challenging to design a valid strategy to compare phylogenetic workflows.

Benchmarking phylogenetic workflows requires knowledge of the true phylogenetic tree, as benchmarking results need to be compared to this reference. The true phylogenetic tree is typically not known in real world settings. As such, various approaches have been proposed to determine or estimate the true phylogenetic tree of a set of strains. Some previous studies have assumed that the consensus of all phylogenies produced by the studied methods is close to the true phylogeny. Alternatively, studies have collated benchmark data sets where the epidemiological data was concordant with the phylogenomic analyses^4^. Because this approach uses real-life data, little is known about the underlying genetic events, and it does not allow to experimentally vary evolutionary parameters. Another approach is to have a mutant strain with an increased mutation rate evolve *in vitro*, and determine the structure of the true phylogeny from the experimental evolution controlled in the lab^5^. This approach provides a good grasp of the true phylogeny and allows the sampling of ancestral strains, but the method is costly and time-consuming and evolutionary parameters cannot be easily controlled. Finally, some studies have used *in silico* evolution to produce realistic sequencing data together with an *a priori* defined true phylogeny^3,6^. This approach offers the possibility to increase or decrease the rate of a range of evolutionary events, such as point mutations, indels, gene duplication, gene loss, gene translocation and lateral gene transfer. Additionally, genomic regions can be evolved under different evolutionary models, as is typical in real life scenarios (e.g. protein-coding genes vs. intergenic regions). Finally, this approach allows a comparison to the true phylogeny, which is not possible with other methods.

In this study, we aim to assess which bioinformatic workflows are able to reconstruct the true phylogeny accurately under diverse evolutionary scenarios. We consider simulating evolution *in silico* to be the optimal approach to achieve this. We simulated the evolution of *Escherichia coli* genomes *in silico* under eight different scenarios, varying the rates of indels, gene duplication, gene loss and lateral gene transfer. We used these simulated datasets to assess the topological accuracy of seventeen phylogenetic reconstruction workflows, including *de novo* genome assembly, alignment or mapping, and finally phylogenetic tree inference. We included six alignment or mapping methods to identify single nucleotide polymorphisms (SNPs) between samples which can be subdivided into k-mer alignment, reference mapping and gene-by-gene alignment methods. We also included three different *de novo* assembly approaches as the impact of this pre-processing step on phylogenomic accuracy is understudied.

## Methods

### Study design

This study consists of two main parts: simulation of *in silico* genome evolution (Fig 1A) and application of phylogenetic workflows on the simulated data sets (Fig 1B). A total of eight sets of parameters were used to simulate a variety of evolutionary processes on genic and intergenic regions separately, using the same phylogeny every time (Table S1). Each simulation was repeated three times with different random seeds to obtain technical replicates. From the *in silico* evolved genomes, short sequencing reads were generated. These sequencing reads were then used as input for the 17 phylogenetic workflows. We tested three *de novo* assembly algorithms in the workflows (Velvet, SKESA, SPAdes), alongside six methods for alignment or mapping (Snippy, Roary, PIRATE, SKA, kSNP, mlst-check). A total of seventeen phylogenetic workflows were tested (Table S2). All phylogenies have been inferred from alignments using IQ-Tree and ModelFinder. As the same phylogeny was used for each simulation, but the parameters for genetic events changed between simulations, each simulated dataset is expected to yield the same genetic distance between isolates (governed by the phylogeny), although the genetic events that have led to this identical genetic distance could be different (governed by the parameters).

**Figure 1.**
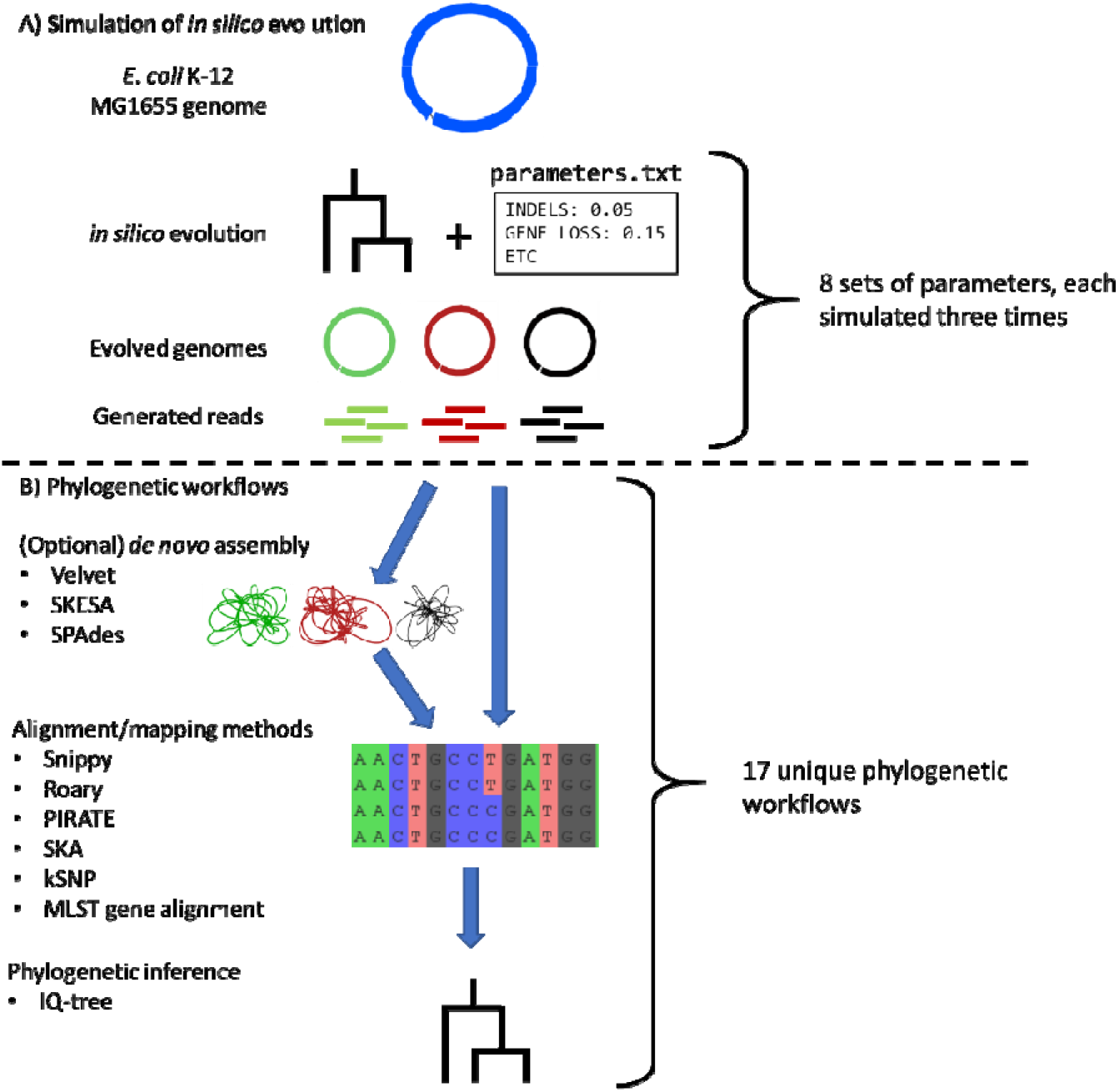
Overview of this study. **A)** Simulation of the in silico evolution. The *E. coli* K-12 MG1655 genome is evolved *in silico* according to a phylogeny (providing genetic distances) and a set of parameters controlling the rates of genetic events (providing which genetic events result in the genetic distance provided by the phylogeny). The resulting genomes are depicted by coloured complete genome graphs visualised in Bandage^34^. The complete genomes are subsequently shredded into sequencing reads. **B)** Phylogenetic workflows. Generated sequencing reads are assembled into draft genomes (coloured draft genome graphs) or directly mapped onto the ancestral genome. From alignments, phylogenetic trees are inferred using IQ-Tree.

### *In silico* evolution

All code is available as a Snakemake v5.8.1^7^ pipeline at https://github.com/niekh-13/phylogenetic_workflows. All tools were run using default parameters, unless otherwise noted. The complete chromosome of *Escherichia coli* K-12 MG1655 (RefSeq Assembly GCF_000005845.2) was used as the ancestral genome in all simulations. Evolution was simulated according to the phylogeny described in Kremer *et al*.^8^. The general approach used in this study was based on the approach described by Lees *et al*.^3^. The ancestral genome was annotated using Prokka v1.14.6^9^ and subsequently divided into protein-coding genes and intergenic regions (all sequences not annotated as protein-coding gene). Protein-coding regions were *in silico* evolved using Artificial Life Framework v1.0 (ALF)^10^ while intergenic regions were *in silico* evolved using DAWG v2.0.beta1^11^.

ALF simulations were run using an empirical codon model, using a standard indel rate of 0.0252, a gene duplication and gene loss rate of 0.05, lateral gene transfer rates of 0.04 for single genes and 0.16 for groups of genes, and no spontaneous gene inversion or gene translocation, based on previous bacterial simulations^3^. Complete specifications for the default run are available from https://github.com/niekh-13/phylogenetic_workflows/blob/master/input/alf_protein_sim.drw. Seven additional simulation were performed (Table S1): “Indel × 0.5” (halved indel rate), “Indel × 2” (doubled indel rate), “gene duplication × 2” (doubled gene duplication rate), “gene loss × 2” (doubled gene loss rate), “gene duplication × 2 & gene loss × 2” (doubled gene duplication and gene loss rates), “lateral gene transfer × 0.5” (halved lateral gene transfer rate for single genes and groups of genes), “lateral gene transfer × 2” (doubled lateral gene transfer rate for single genes and groups of genes).

DAWG simulations were run using a default indel rate of 0.00175 and evolved under a general time-reversible (GTR) model with rates A⟷C: 0.91770, A⟷G:4.47316, A⟷T:1.10375, C⟷G:0.56499, C⟷T:6.01846, G⟷T:1.00000, based on the GTR matrix inferred from dataset of nearly 1200 *E. coli* strains isolated from various host species (HECTOR study, manuscript in preparation). For simulations “indel × 0.5” and “indel × 2” the indel rate was appropriately changed (Table S1).

Per simulation, ALF and DAWG *in silico* evolution yielded protein-coding genes and intergenic regions for 96 *in silico* evolved genomes. These were assembled into 96 complete genomes. As stop codons are removed during ALF simulation, stop codons were inserted at the ends of genes. Illumina sequencing reads were simulated using ART v2016.06.05^12^, using seed 21.

### Comparing pipelines

From the simulated Illumina sequencing reads, phylogenies were reconstructed through seventeen workflows (Table S2). Assemblies were created using the Shovill v1.1.0 (https://github.com/tseemann/shovill) wrapper for Velvet v1.2.10^13^, SPAdes v3.14.0 using “-- isolate” mode^14^ and SKESA v2.3.0^15^. Contigs were retained if they were 500bp or larger for all *de novo* assembly algorithms. Assembly quality metrics were assessed using Quast v5.0.2^16^ and all versus all average nucleotide identity (ANI) comparisons were made using fastANI v1.2^17^. K-mer alignment methods kSNP v3.1^18^ and SKA v1.0^19^ were used on all assemblies, and SKA was additionally run on sequencing reads. In our study, both tools were used to extract k-mers of 31 bp from assemblies or sequencing reads. Subsequently, these tools aligned k-mers of which the first and last 15 bp were identical, thus allowing only the middle base to vary between aligned k-mers. This k-mer alignment produced SNP alignments, which can be used for phylogenetic inference. Important to note is that although SKA and kSNP also employ k-mer-based methods, these methods are conceptually distinct from other k-mer-based tools such as Mash (https://github.com/marbl/Mash). The bacterial mapping pipeline Snippy v4.6.0 (https://github.com/tseemann/snippy) was used on sequencing reads alone. Gene-by-gene methods Roary v3.13.0^20^ and PIRATE v1.0.3^21^ were used on annotations produced by Prokka v1.14.6. Finally, alignments were constructed from multi-locus sequence type genes using mlst-check v2.1.1706216^22^ and realigned using ClustalO v1.2.4^23^. All methods, including k-mer alignment methods, produce nucleotide alignments, which were subsequently used to infer phylogenies using IQ-tree v2.0.3^24^ and ModelFinder^25^ packaged with IQ-tree. Differences between the ground truth phylogeny and produced phylogenies were assessed using Robinson-Foulds distance calculation implemented in ape v5.4^26^ and Kendall-Colijn distance calculation implemented in treespace v1.1.3.2^27^. All simulations and pipelines were run three times, with seeds 1, 42 and 1704 in ALF simulation. Alignment lengths were extracted using snp-sites v2.5.1 (REF).

### Visual and statistical analysis

Parsing of results was performed using the pandas library v0.25.3^28^ in Python v3.8.3 and using the tidyverse v1.3.0^29^ and rstatix v0.6.0 (https://cran.r-project.org/package=rstatix) libraries in R v4.0.1. Results were plotted using ggplot2 v3.3.1^30^, ggpubr v0.4.0 (https://cran.r-project.org/package=ggpubr), ggthemes v4.2.0^31^, patchwork v1.0.1^32^ and using SuperPlotsOfData^33^. Tests for statistical significance were carried out in SuperPlotsOfData using paired Welch’s t-tests where indicated. Bonferroni correction for multiple testing was applied where applicable.

## Results

### Reference-based mapping and k-mer alignment methods yield phylogenetic trees most similar to ground truth

The *in silico* evolution yielded isolate sets with a genetic diversity comparable to a single bacterial species (≥95% average nucleotide identity^35^, Figures 1 and S1). The same level of genetic diversity was attained between simulations, although these simulations included different rates of simulated genetic events (substitutions, indels, lateral gene transfer, etc., Table S3).

The optimal phylogenetic workflow should produce a phylogeny identical to the phylogeny which was used in the simulation process (the “ground truth phylogeny”). Per workflow, we calculated tree distance between the phylogeny produced by the workflow and the ground truth phylogeny. Tree distances were expressed in the Robinson-Foulds distance and the Kendall-Colijn metric.

The workflow showing the lowest tree distances across simulations employed SPAdes *de novo* assembly and subsequently SKA for k-mer alignment. After Bonferroni correction for multiple testing, the Kendall-Colijn metric of this workflow was significantly lower than all other workflows except Snippy, SPAdes + kSNP, SKESA + kSNP and SKESA + SKA (Fig 2 and Table S4). Notably, core gene alignment methods and methods employing Velvet for *de novo* assembly performed worse in our study. MLST gene alignment methods showed the highest deviation from the ground truth phylogeny as measured by Kendall Colijn metric and Robinson Foulds distance (Fig S2).

**Figure 2.**
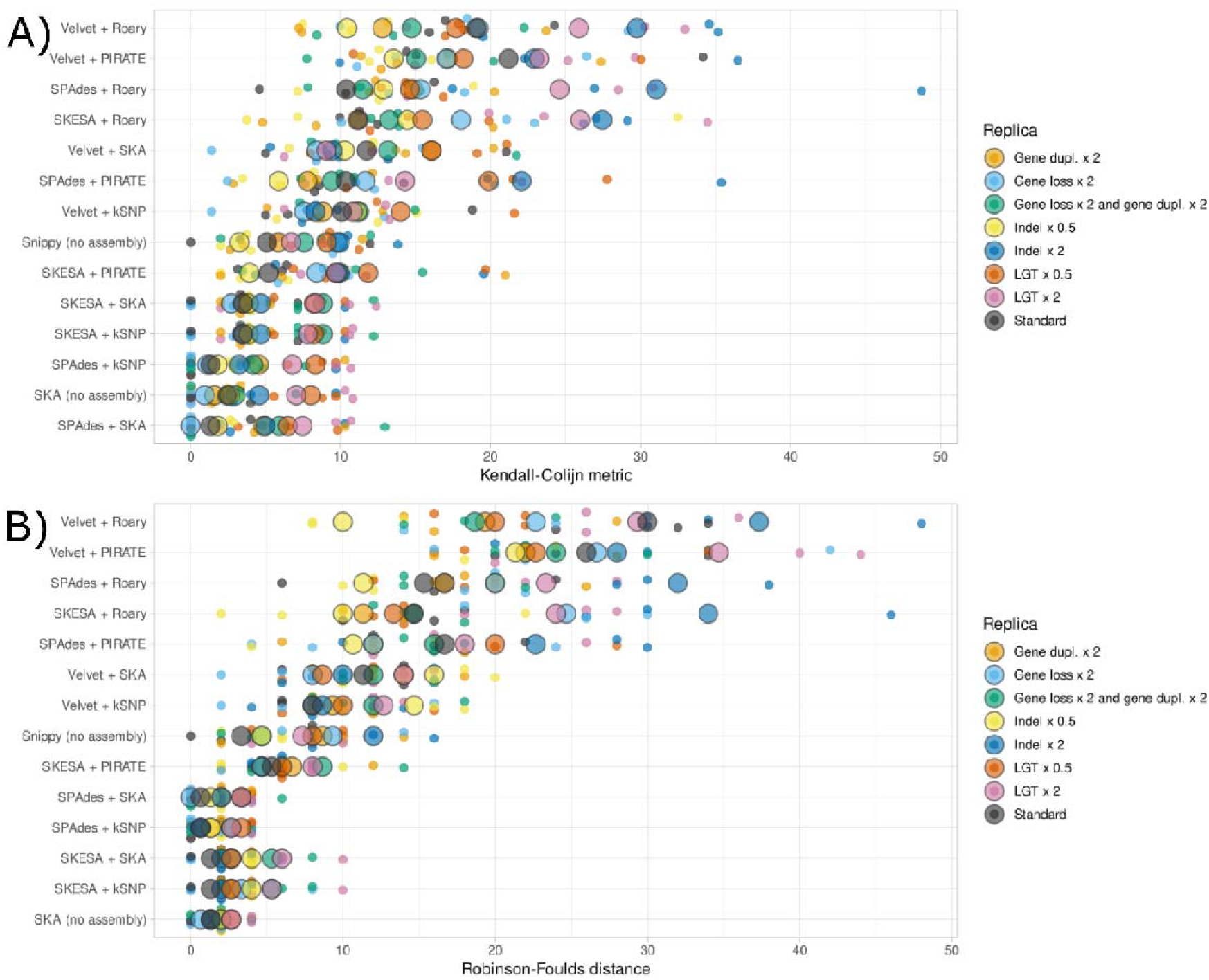
Kendall-Colijn metrics and Robinson-Foulds distances per phylogenetic workflow across eight simulations. Displayed distances are calculated between the ground truth phylogeny and the phylogeny produced by the relevant workflow. Generated using SuperPlotsOfData, ordered by median. Large circles indicate median of replica. Small circles indicate separate measurements for replica.

### *De novo* assembly algorithms have a strong influence on accuracy of phylogenetic reconstruction

Next, we compared the accuracy of phylogenetic reconstruction between workflows employing different *de novo* assembly algorithms (Fig 3 and Table S5). Across eight simulations, workflows employing SPAdes and SKESA both resulted in significantly lower Kendall-Colijn metric values compared to the same workflows employing Velvet. In other words, workflows employing SPAdes and SKESA reconstruct phylogenies more accurately than the same workflows employing Velvet.

**Figure 3.**
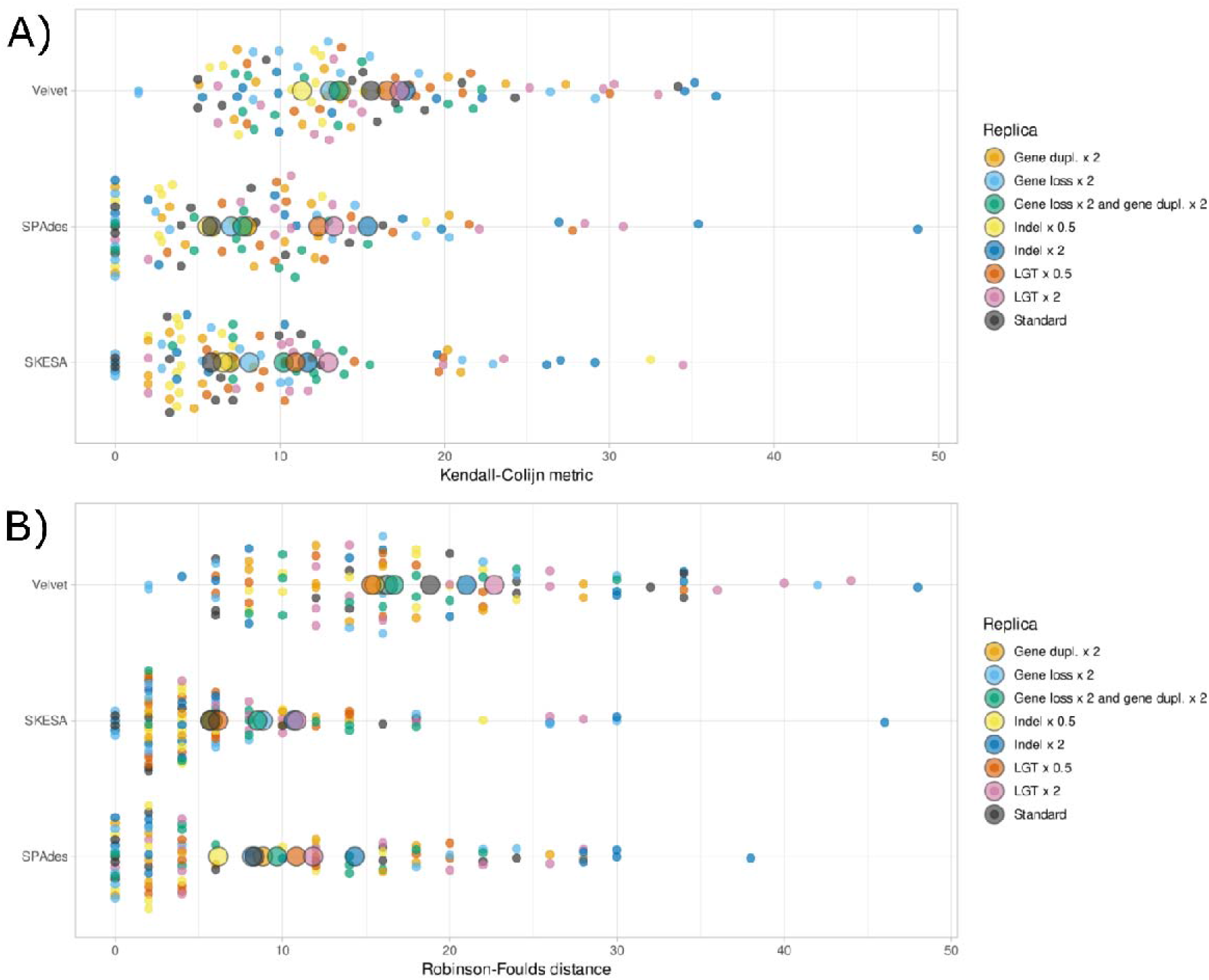
Kendall-Colijn metrics and Robinson-Foulds distances per *de novo* assembly algorithm used in workflows, across eight simulations. Displayed distances are calculated between the ground truth phylogeny and the phylogeny produced by the relevant workflow. Generated using SuperPlotsOfData, ordered alphabetically. Large circles indicate median of replica. Small circles indicate separate measurements for replica.

To gain insights in the *de novo* genome assembly quality, we compared the assemblies produced by Velvet, SKESA and SPAdes to the *in silico* evolved genomes from which the sequencing reads were generated, using detailed assembly quality metrics such as total genome fraction, NGA50, and the number of misassemblies alongside standard quality metrics such as number of contigs or total assembly size. We observed that although Velvet produced genome assemblies with a relatively high NGA50, Velvet also produced the highest number of misassemblies compared to SKESA or SPAdes (Table S6 and Fig S3). SPAdes seemed to perform best across multiple assembly quality metrics, reconstructing a large part of the original genome in few contigs (NGA50, genome fraction reconstructed, number of contigs), with a low number of errors (number of misassemblies).

### Accuracy of phylogenetic reconstruction is associated with number of informative sites in the alignment

We hypothesised that the workflows using a larger part of the genome in the comparative analysis would yield larger alignments and more accurate phylogenetic reconstruction. To assess this, we extracted the alignment length produced per workflow. We found that the alignment length shows a strong negative correlation with the Kendall-Colijn metric and explains approximately 32% of variance in KC metric (R^2^, Fig 4). This indicates that the methods that included a larger fraction of the genomes under study produced more accurate phylogenies. When the workflows employing MLST alignments were included, this negative correlation was even stronger (Fig S4).

**Figure 4.**
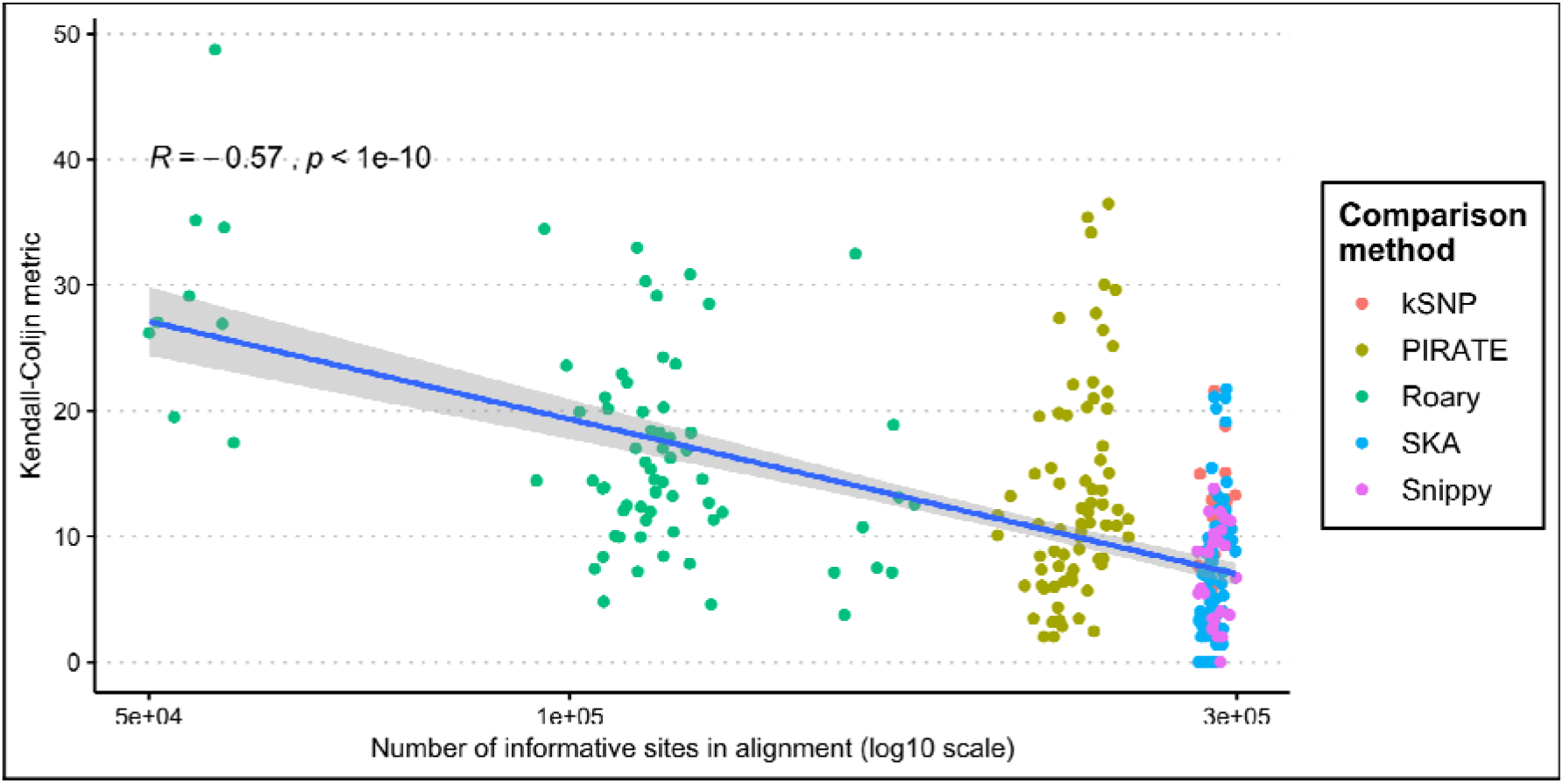
Count of informative sites in alignment plotted against Kendall-Colijn metric, with a linear model fitted (shading indicates 95% confidence interval). Pearson’s Rho and associated p-value are shown.

### Phylogenetic benchmarking shows a high variability between replicates

Repeating each of the eight simulations three times allows us to assess the reproducibility of this analysis. We see extensive variability in the accuracy of phylogenetic reconstruction even when comparing identical workflows across identical simulations, where only the starting seed for simulation differed (Fig 5). The largest difference between technical replicates reached a 31 point different in the Kendall-Colijn metric (SPAdes + Roary, simulation with double indel rate). Over 22% of Kendall-Colijn metric calculations were off more than 10 points between technical replicates.

**Figure 5.**
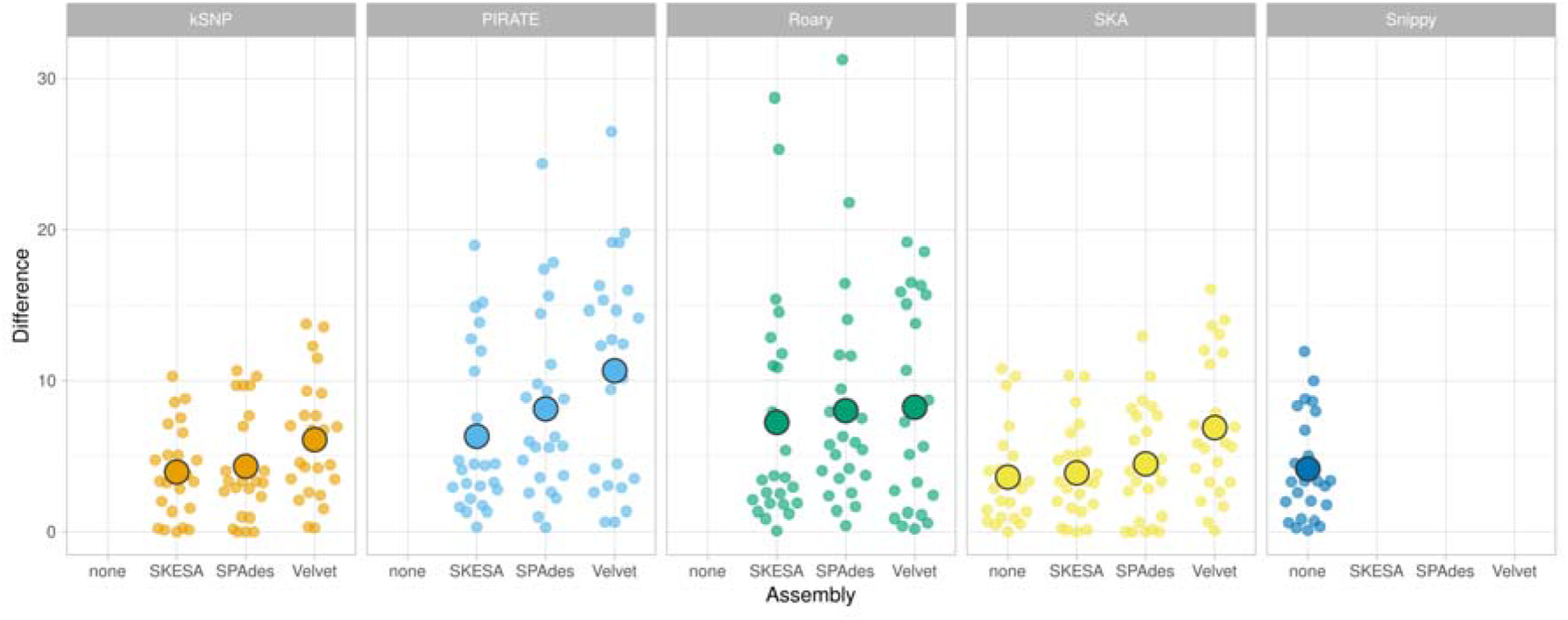
Differences between technical replicates for identical workflows across identical simulations, only differing in starting seed for the simulation. Workflows including MLST were excluded. Generated using SuperPlotsOfData.

## Discussion

We present a systematic analysis of the accuracy of phylogenetic reconstruction of several workflows, based on simulated bacterial whole-genome data. We have included seventeen phylogenetic workflows. These were each benchmarked using eight simulation scenarios with three independent replicates.

First, we show that k-mer alignment methods provide a good alternative to reference-based mapping in species-level phylogenetic reconstruction. The high accuracy of workflows employing k-mer alignment seems to be due to the large fraction of genomes that can be utilized in these workflows, reflected by the high number of informative sites in alignments produced by k-mer methods. Through including eight simulation scenarios we were able to determine a clear influence of the *de novo* assembly algorithm on phylogenetic accuracy. Based on assembly quality evaluation, we hypothesise that an increased rate of misassemblies has a detrimental effect on phylogenetic accuracy. This also applies to k-mer alignment methods, which performed best when combined with either SPAdes or SKESA.

Surprisingly, we observed a high variability between replicates of phylogenetic workflows. Over one-fifth of comparisons showed differences of 10 points or more in the Kendall-Colijn metric. To contextualize, the difference in median Kendall-Colijn metric between the best and worst workflows in Fig 2A was 14.6 points. Generally, workflows using core gene alignment methods such as Roary or PIRATE displayed the highest discrepancies between replicates. This might be because core gene alignment methods need to employ heuristics to compare genes in an all-versus-all manner, which could introduce variability in their results.

Across seventeen phylogenetic workflows, eight simulations and three replicates, we reconstructed a total of 408 phylogenies. By including multiple workflows, simulations and replicates, this number increases quickly. We were able to limit computational workload by selecting only a single method (IQtree) to infer phylogenies from alignments. We chose to include only IQ-Tree because there was little difference between IQ-Tree, RAxML or other approaches in earlier studies^3^, because IQ-Tree is widely used and thus represents an established method to infer phylogenies, and finally because IQ-Tree offers the identification of an optimal substitution model through ModelFinder.

One of the challenges in benchmarking studies is to employ all methods in such a way that these can be compared sensibly. For k-mer alignment methods SKA and kSNP, we observed that configuring the desired k-mer length differs between tools. To obtain aligned k-mers of 31 bp, SKA requires to set k-mer length (flag “-k”) to 15, resulting in the alignment of two split k-mers of 15 bp with a middle base, amounting to a total aligned k-mer of 31 bp.

However, for kSNP the k-mer length (flag “”) should be set to 31, to obtain a 31 bp aligned k-mers of which the middle base may vary. Configuring the k-mer length correctly resulted in a highly similar accuracy of SKA and kSNP, while previous studies did not establish similar performance due to discrepancies in k-mer length configuration^19^.

Determining the exact rates of genetic events such as point mutations or indels is challenging. In this study, we have evolved bacterial genomes across a range of evolutionary scenarios which means our results should be interpreted as generalisable findings, rather than findings specific to *Escherichia coli* and its evolutionary mechanisms.

Here we simulated datasets which exhibited a limited genetic diversity, similar to the genetic diversity observed within species (at least ~95% ANI^35^). In the context of more diverse datasets, for example comparing different species or genera, we expect that k-mer alignment methods would perform worse as these methods typically perform best with limited genetic diversity^19^. In accord with our results, we theorize that this is due to a faster decrease in informative sites with increasing evolutionary distance.

This study illustrates how phylogenetic reconstruction methods based on bacterial whole genome data compare. The simulations cover diverse evolutionary scenarios for bacterial species, providing detailed insight into the performance of phylogenetic reconstruction methods valid across diverse sets of bacterial strains. Recently developed k-mer alignment methods achieved similar accuracy as the gold standard (reference mapping) and thus seems to be a useful alternative when no suitable reference genome is available. Every microbe evolves according to different evolutionary parameters, so phylogenetic workflows need to be able to resolve many different evolutionary scenarios. Our study provides data on the accuracy of existing phylogenetic workflows and a framework to assess future phylogenetic workflows.

## Author statements

### Authors and contributors

Conceptualization: BCLP, DRM, CS. Data curation: BCLP, NAHH. Formal Analysis: BCLP, NAHH. Funding acquisition: CS. Investigation: BCLP, NAHH. Methodology: BCLP, NAHH, DRM. Project administration: BCLP, DRM, CS. Software: BCLP, NAHH. Supervision: DRM, CS. Validation: BCLP, NAHH. Visualization: BCLP, NAHH. Writing – original draft: BCLP, NAHH. Writing – review & editing: BCLP, NAHH, DRM, CS.

## Acknowledgements

We thank SURFsara (www.surfsara.nl) for the support in using the Lisa Compute Cluster.

We thank the members of the HECTOR consortium for the use of the HECTOR data in inferring the GTR matrix used in the *in silico* evolution.

## Supplement

**Fig S1.**
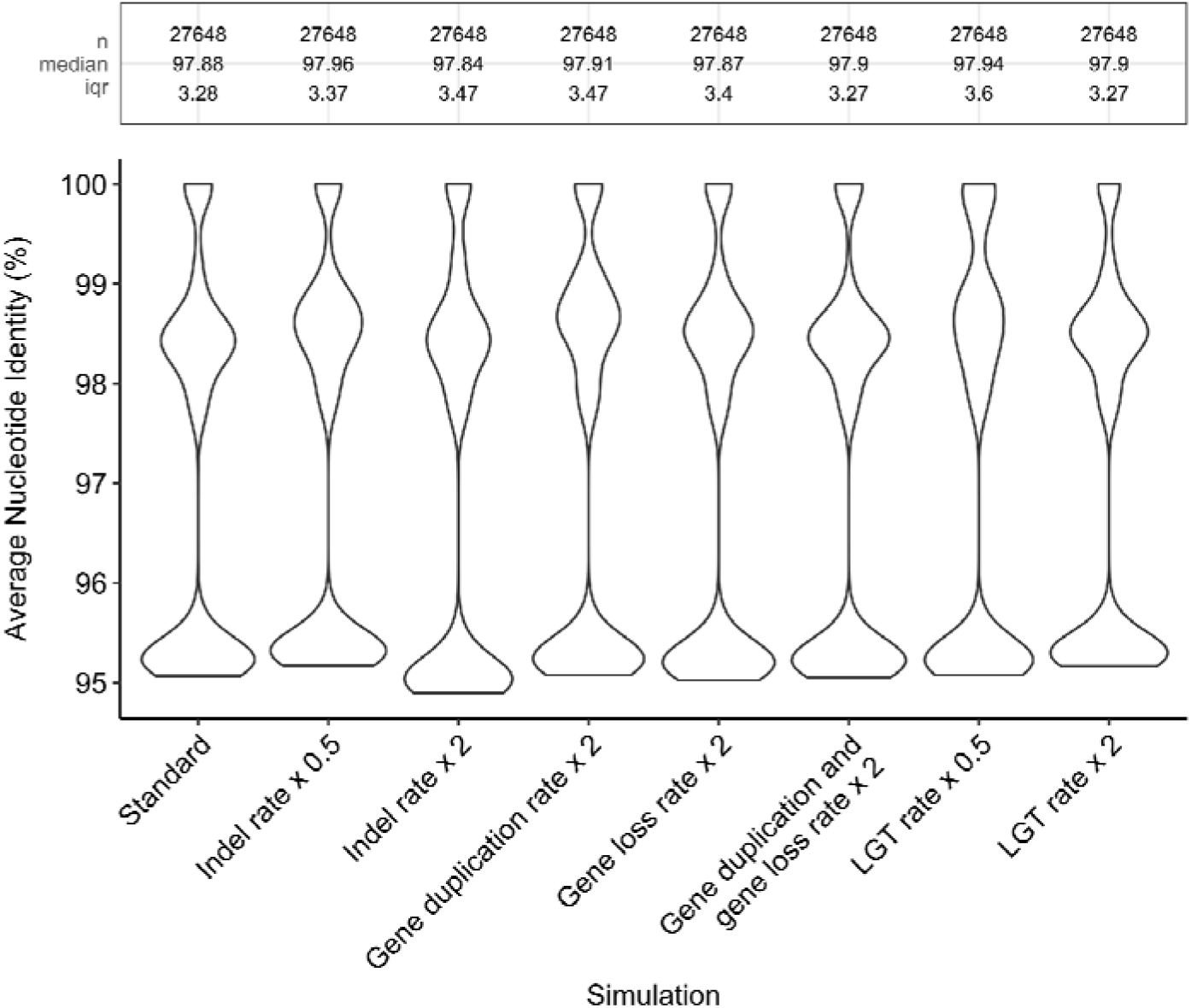
Violin plots of ANI comparisons made using FastANI, per simulation. From each simulation replicate (three replicates for eight simulations), all 96 *in silico* evolved genomes were compared in an all vs. all fashion.

**Figure S2.**
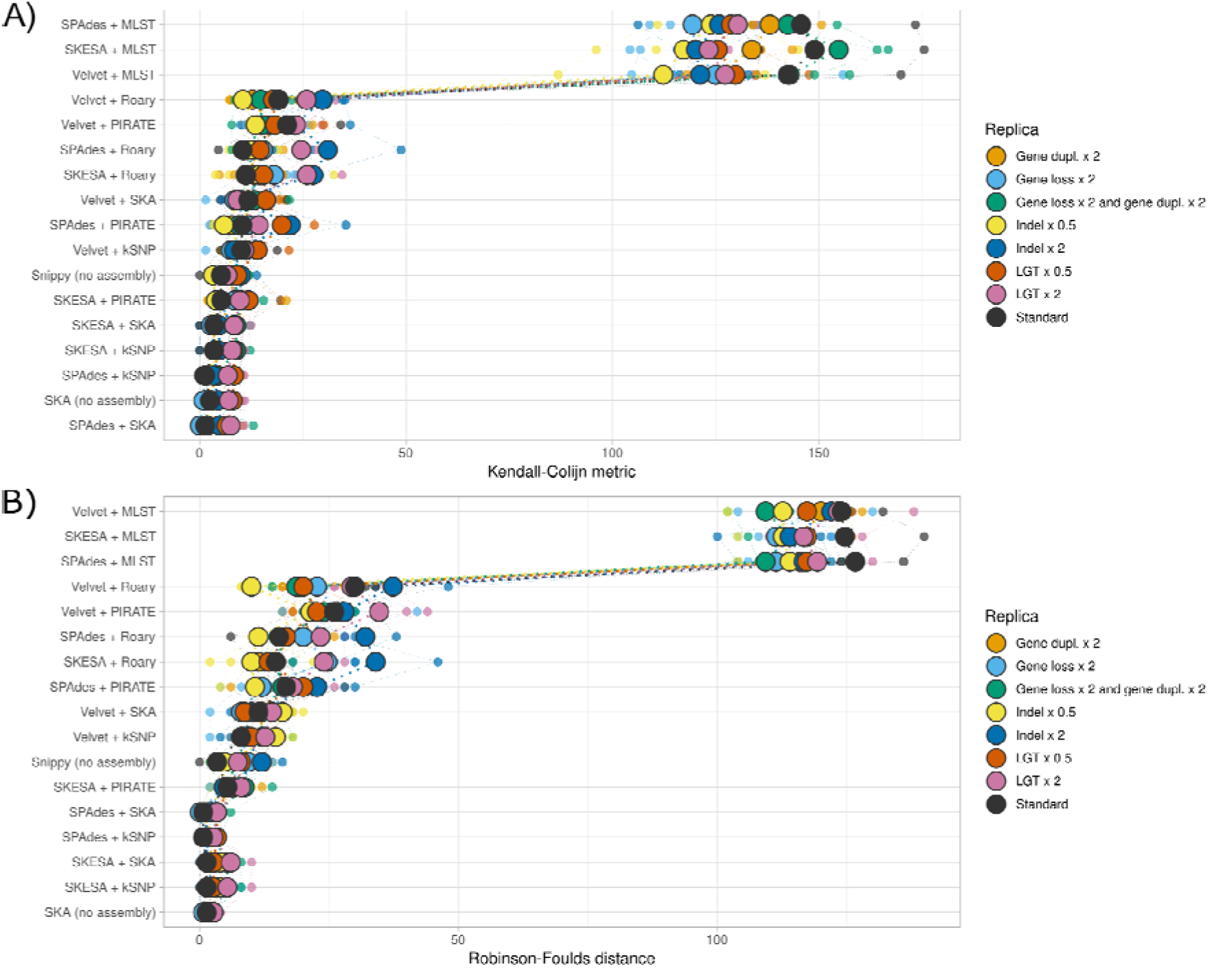
Kendall-Colijn metrics and Robinson-Foulds distances between the ground truth phylogeny and phylogenies produced by workflows, across eight simulations.

**Figure S3.**
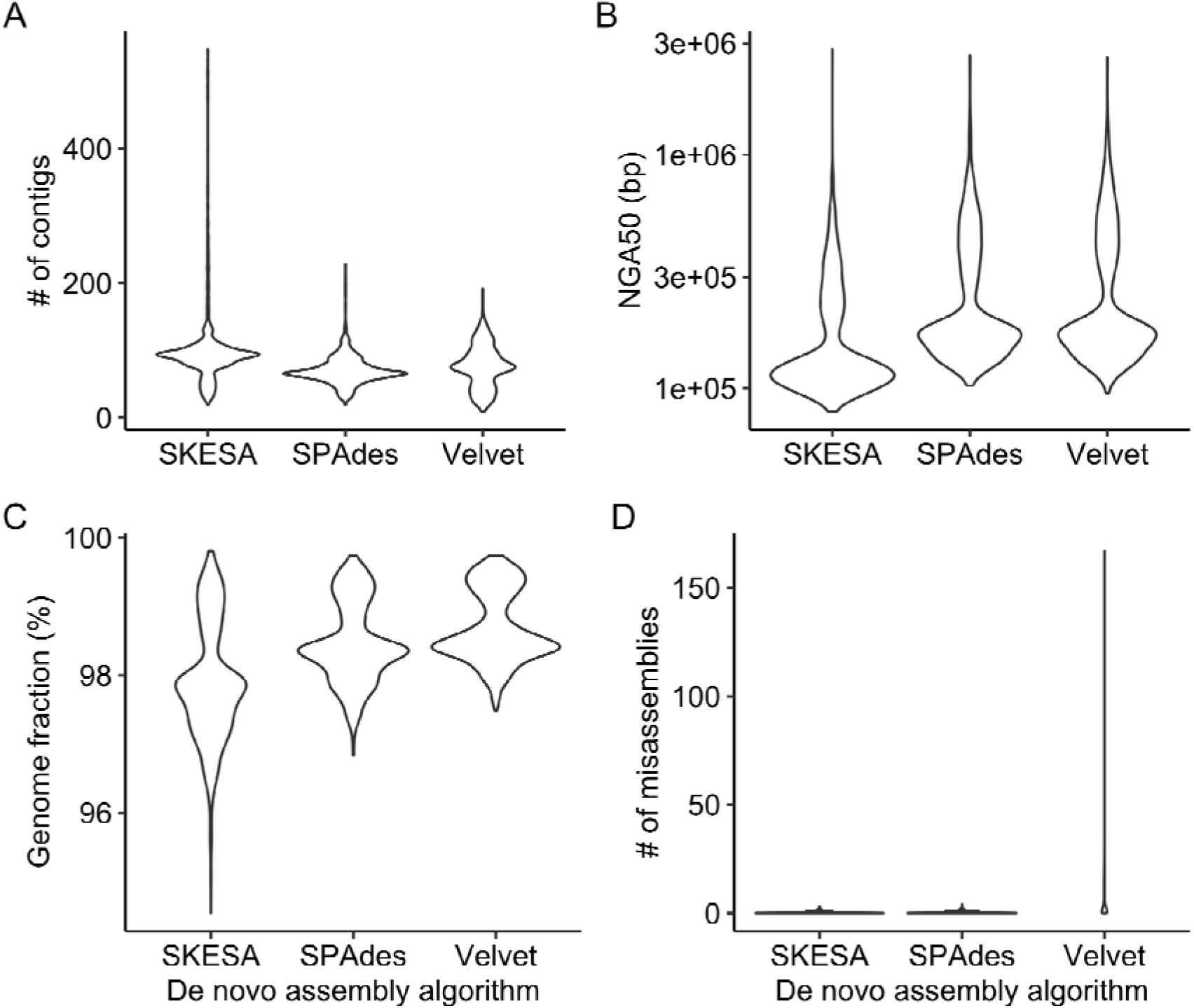
Comparison of SKESA, SPAdes and Velvet algorithms for *de novo* genome assembly, based on number of contigs, NGA50, genome fraction reconstructed and number of misassemblies.

**Figure S4.**
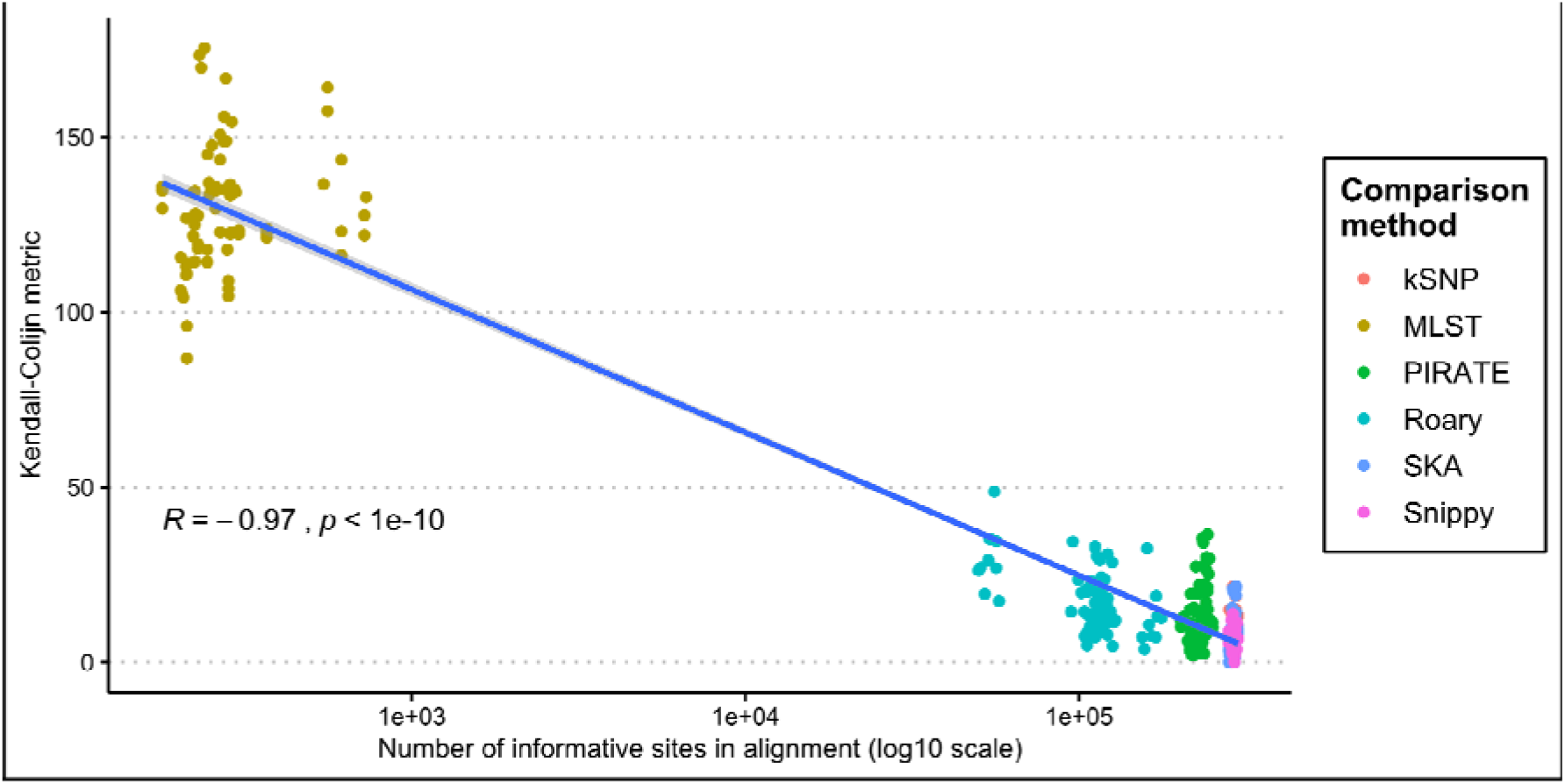
Count of informative sites in alignment plotter against Kendall-Colijn metric, with a linear model fitted (shading indicates 95% confidence interval). Pearson’s Rho and associated p-value are shown.

**Table S1.** Parameters which differed per simulation. All other parameters were kept stable.

| Simulation | Genes (ALF) |  |  |  |  | Intergenic regions<br>(DAWG) |
| --- | --- | --- | --- | --- | --- | --- |
|  | Indel rate | geneDuplRate | geneLossRate | lgtRate | lgtGRate | Indel rate |
| Standard | 0.0252 | 0.05 | 0.05 | 0.04 | 0.16 | 0.001750 |
| Indel x 0.5 | 0.0126 | 0.05 | 0.05 | 0.04 | 0.16 | 0.000875 |
| Indel x 2 | 0.0504 | 0.05 | 0.05 | 0.04 | 0.16 | 0.003500 |
| LGT x 0.5 | 0.0252 | 0.05 | 0.05 | 0.02 | 0.08 | 0.001750 |
| LGT x 2 | 0.0252 | 0.05 | 0.05 | 0.08 | 0.32 | 0.001750 |
| Gene dupl. x 2 | 0.0252 | 0.10 | 0.05 | 0.04 | 0.16 | 0.001750 |
| Gene loss x 2 | 0.0252 | 0.05 | 0.10 | 0.04 | 0.16 | 0.001750 |
| Gene loss x 2<br>and gene dupl. x<br>2 | 0.0252 | 0.10 | 0.10 | 0.04 | 0.16 | 0.001750 |

**Table S2.**
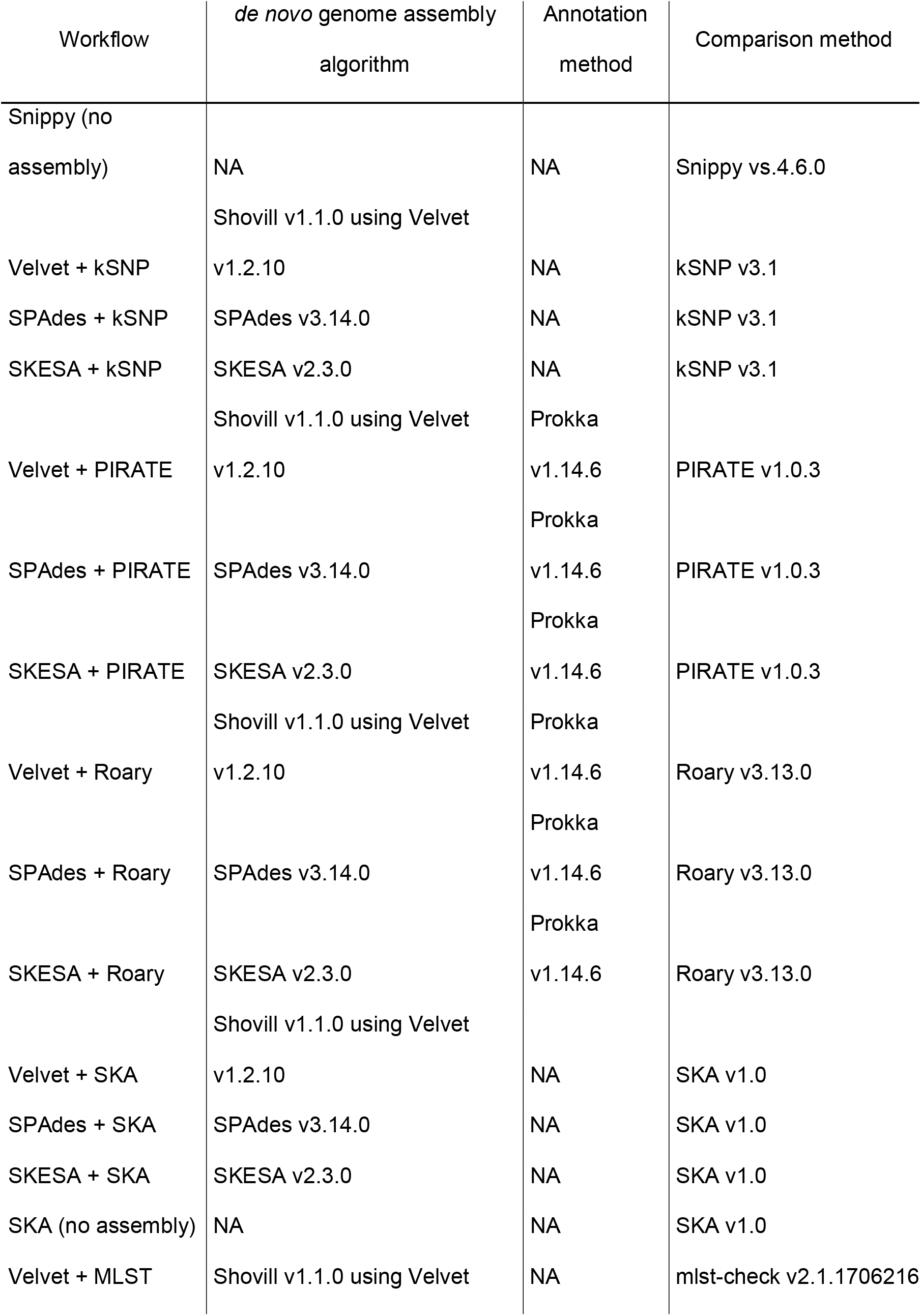
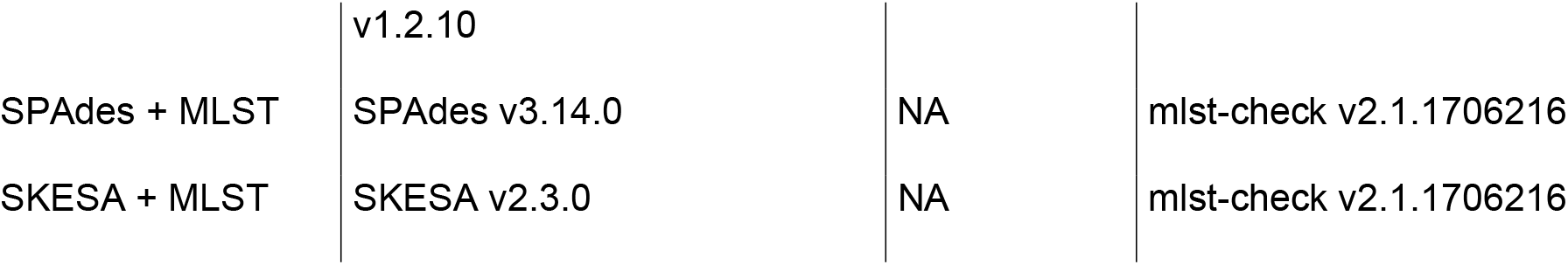
Seventeen phylogenetic workflows included in this study, with software versions.

**Table S3.**
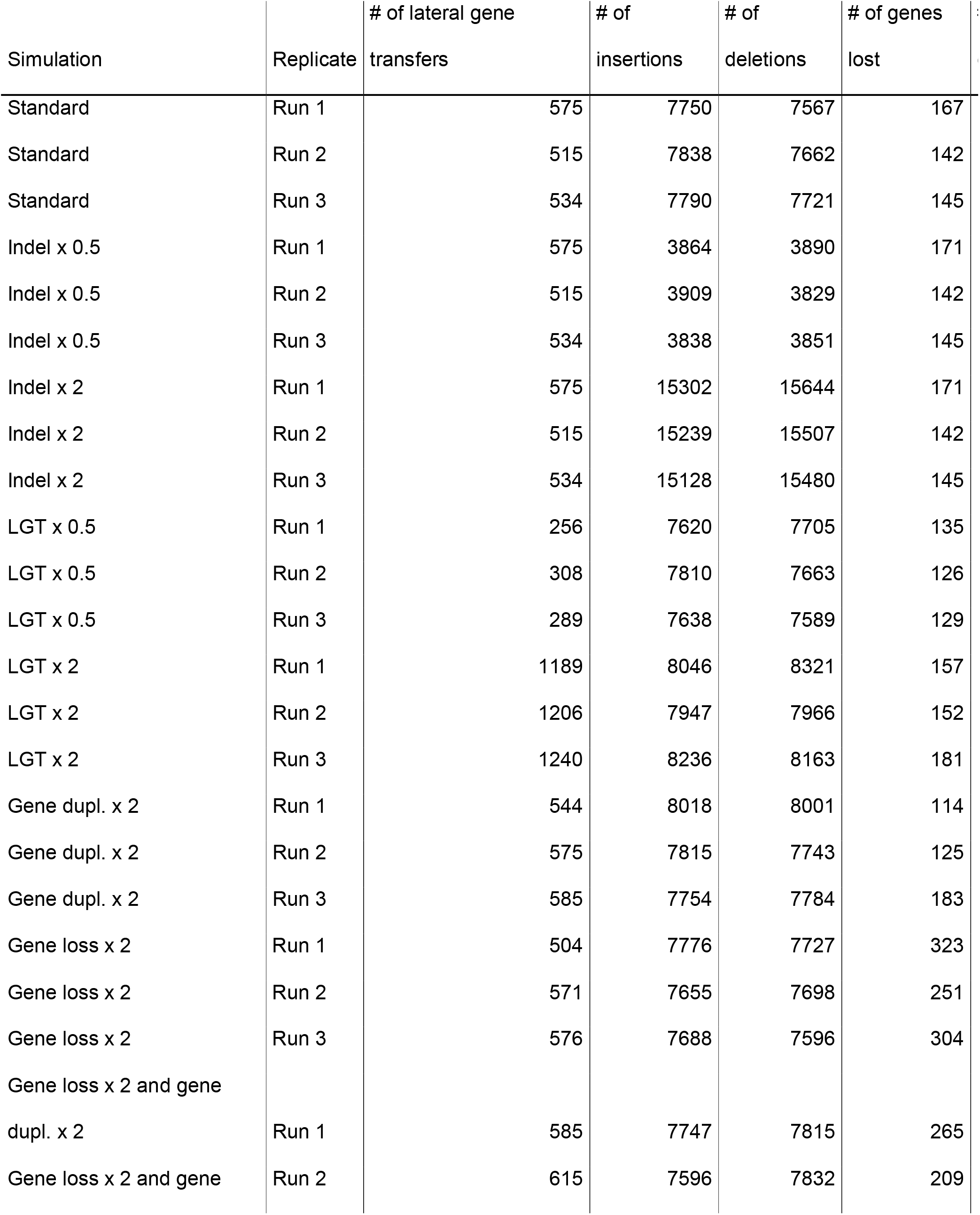
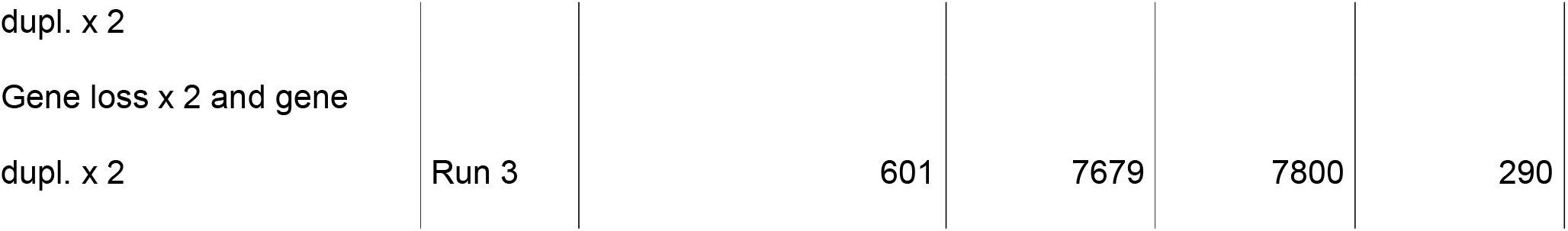
Counts of genetic events across replicates of eight simulations.

**Table S4.** Results of statistical significance tests using Welch’s t-test between SPADES + SKA vs other workflows. The p-value threshold after Bonferroni correction for multiple testing is 3.85×10^−3^. Statistically significant differences are marked with an asterisk.

| Condition | Difference<br>(95% confidence<br>interval) | P-value |
| --- | --- | --- |
| SPAdes + SKA vs Velvet +<br>PIRATE | 14.43 (11.30 to 17.57) | 0.0000121* |
| SPAdes + SKA vs Velvet + kSNP | 6.14 (4.11 to 8.18) | 0.0001875* |
| SPAdes + SKA vs Velvet + SKA | 7.67 (4.98 to 10.36) | 0.0002644* |
| SPAdes + SKA vs Velvet + Roary | 14.59 (9.36 to 19.82) | 0.0003062* |
| SPAdes + SKA vs SKESA +<br>PIRATE | 4.47 (2.81 to 6.13) | 0.0003734* |
| SPAdes + SKA vs SKESA + Roary | 13.01 (8.03 to 18.00) | 0.0004572* |
| SPAdes + SKA vs SPAdes +<br>Roary | 12.79 (7.32 to 18.27) | 0.0008835* |
| SPAdes + SKA vs SPAdes +<br>PIRATE | 8.56 (4.24 to 12.88) | 0.0022377* |
| SPAdes + SKA vs Snippy (no<br>assembly) | 3.05 (0.36 to 5.74) | 0.0316289 |
| SPAdes + SKA vs SKESA + SKA | 1.35 (0.10 to 2.60) | 0.0375962 |
| SPAdes + SKA vs SKESA + kSNP | 1.38 (0.00 to 2.76) | 0.0496085 |
| SPAdes + SKA vs SKA (no<br>assembly) | -0.33 (-1.87 to 1.21) | 0.6238804 |
| SPAdes + SKA vs SPAdes + kSNP | -0.19 (-1.24 to 0.86) | 0.6774758 |

**Table S5.** Results of statistical significance tests using paired Welch’s t-tests between workflows employing Velvet vs SPAdes and SKESA. The p-value threshold after Bonferroni correction for multiple testing is 2.5×10^−2^. Statistically significant differences are marked with an asterisk.

| Condition | Difference (95% confidence interval) | P-value |
| --- | --- | --- |
| Velvet vs SKESA | -5.66 (-7.26 to -4.05) | $7.09 \times 10^{-5*}$ |
| Velvet vs SPAdes | -5.42 (-7.21 to -3.62) | $1.87 \times 10^{-4*}$ |

**Table S6.**
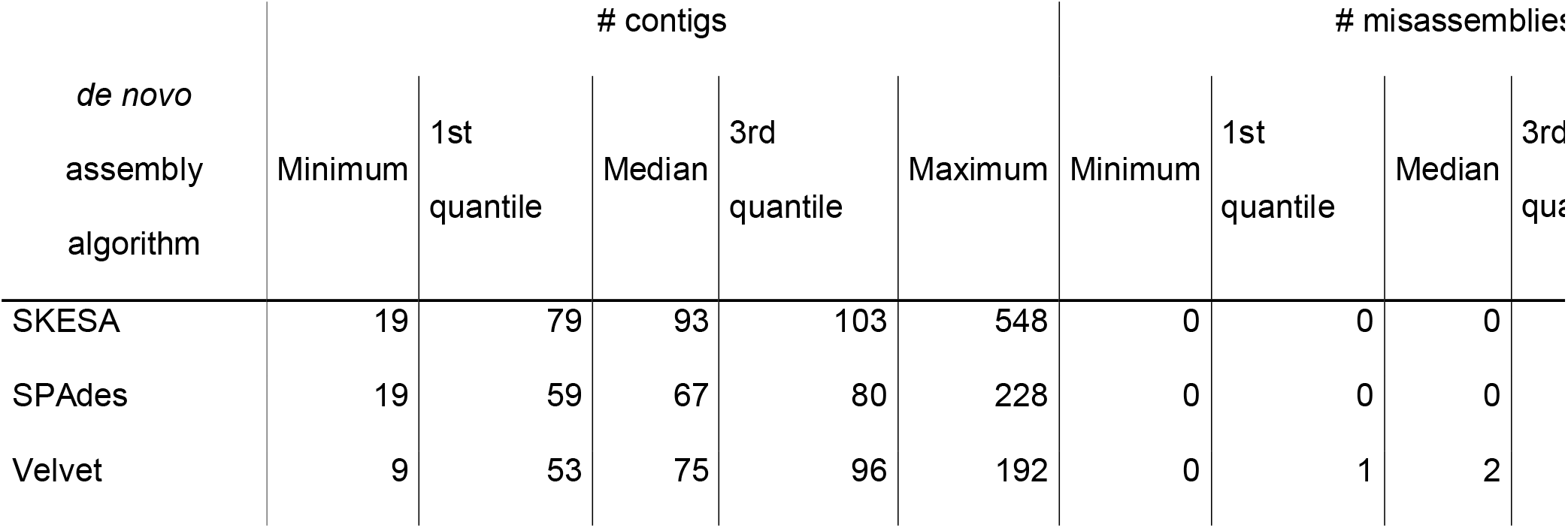
Quast evaluation of *de novo* genome assemblies produced by SKESA, SPAdes and Velvet.

