## Supplementary figures and images for "Benchmarking topological accuracy of bacterial phylogenomic workflows using *in silico* evolution"

### Figure S1

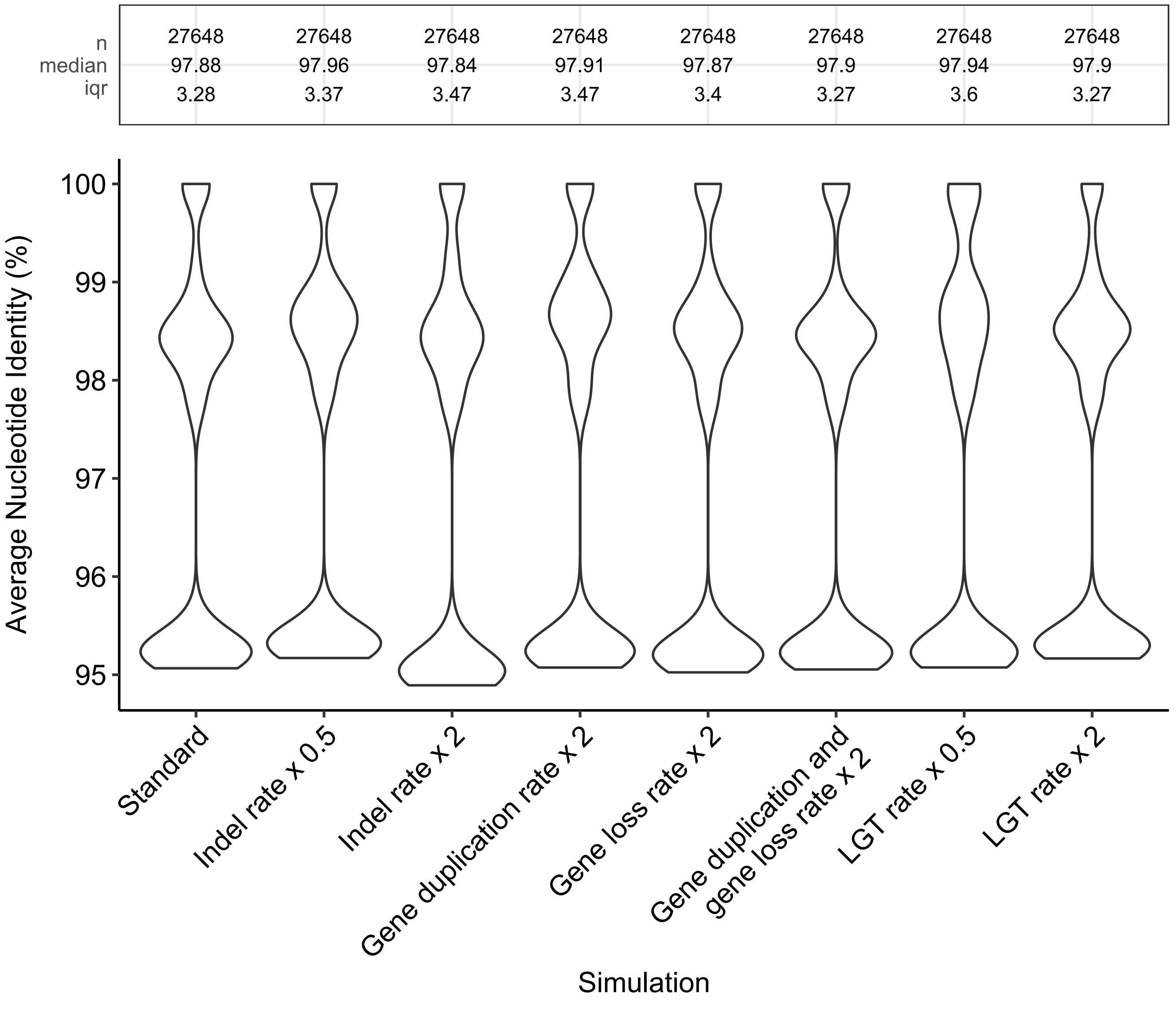

### Figure S2

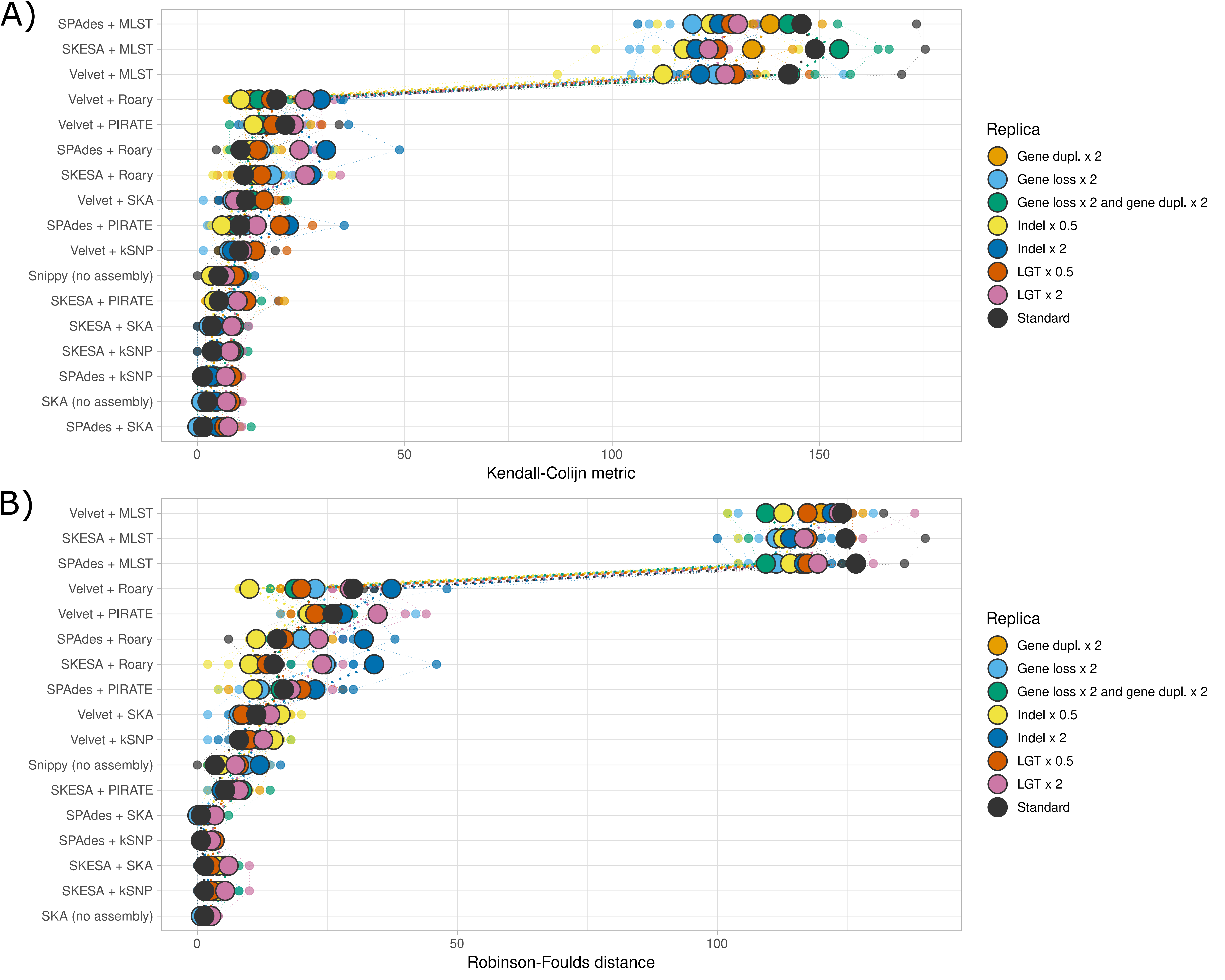

### Figure S3

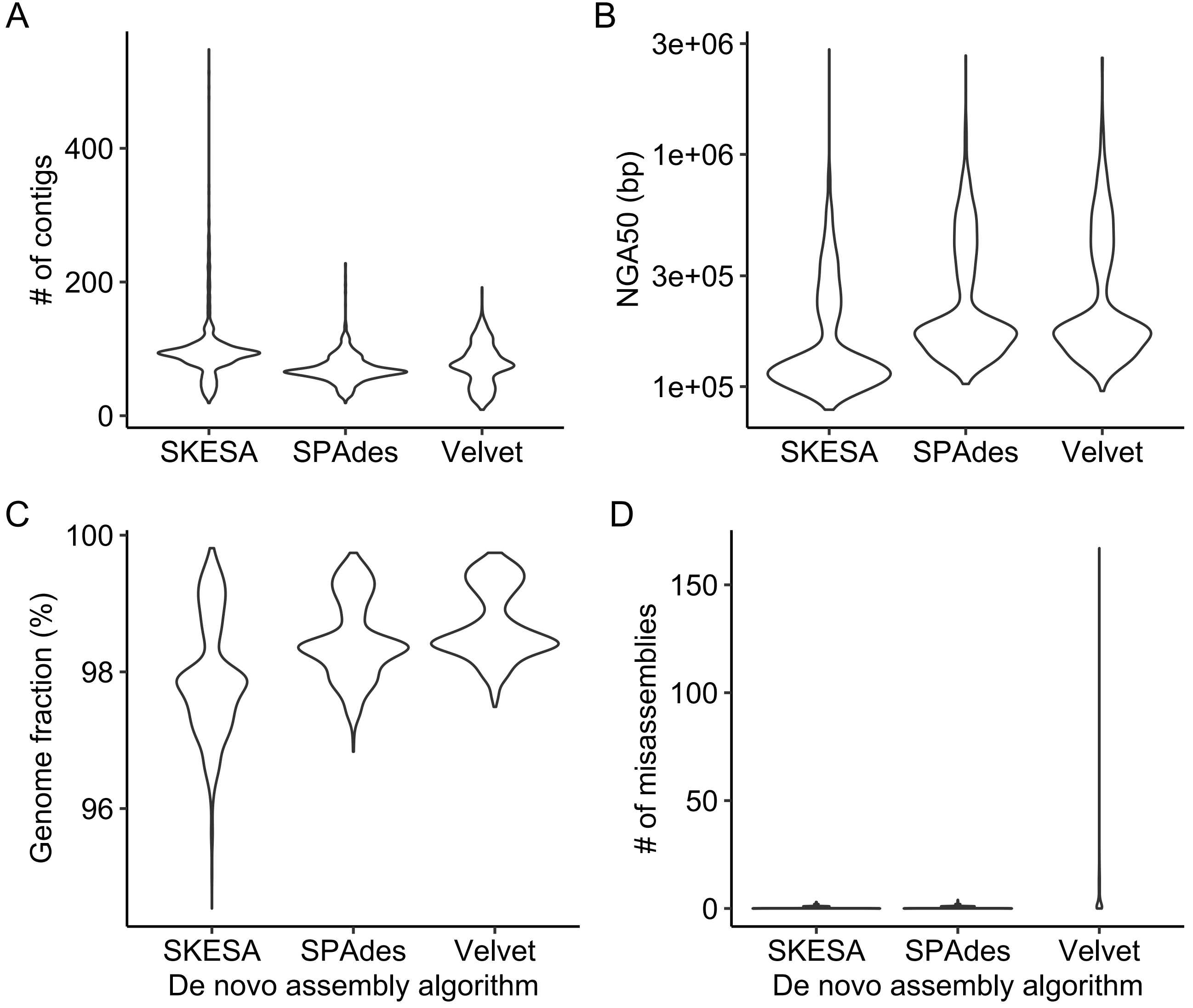

### Figure S4

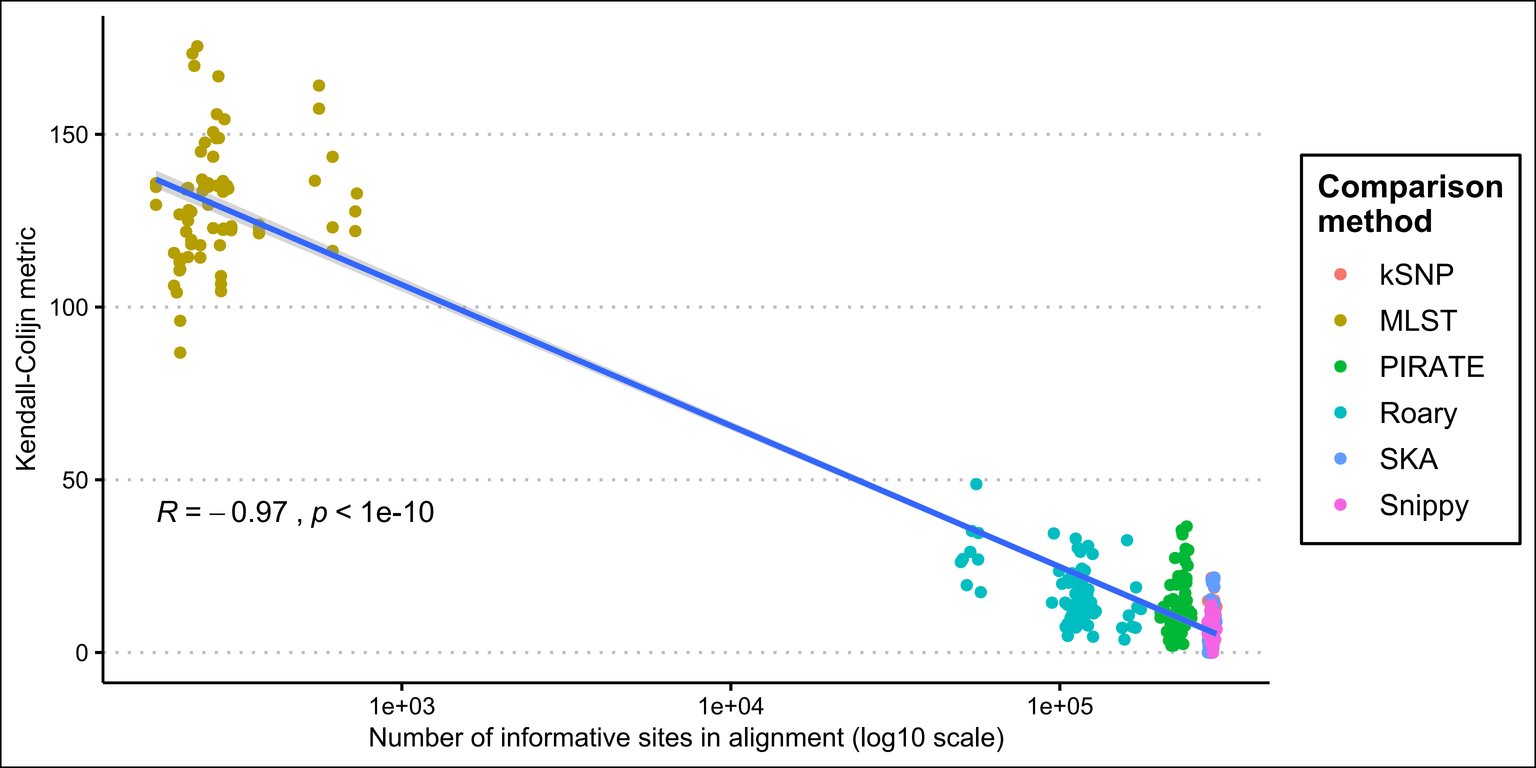
